# Estimating Genome-wide Phylogenies Using Probabilistic Topic Modeling

**DOI:** 10.1101/2023.12.20.572577

**Authors:** Marzieh Khodaei, Scott V. Edwards, Peter Beerli

## Abstract

Methods for rapidly inferring the evolutionary history of species or populations with genome-wide data are progressing, but computational constraints still limit our abilities in this area. We developed an alignment-free method to infer genome-wide phylogenies and implemented it in the Python package TopicContml. The method uses probabilistic topic modeling (specifically, Latent Dirichlet Allocation or LDA) to extract ‘topic’ frequencies from *k*-mers, which are derived from multilocus DNA sequences. These extracted frequencies then serve as an input for the program Contml in the PHYLIP package, which is used to generate a species tree. We evaluated the performance of TopicContml on simulated datasets with gaps and three biological datasets: (1) 14 DNA sequence loci from two Australian bird species distributed across nine populations, (2) 5162 loci from 80 mammal species, and (3) raw, unaligned, non-orthologous PacBio sequences from 12 bird species. Our empirical results and simulated data suggest that our method is efficient and statistically robust. We also assessed the uncertainty of the estimated relationships among clades using a bootstrap procedure.

## Introduction

Phylogenetic analysis traditionally relies on the alignment of orthologous sequence data, a process that can be challenging due to the complexity of genomic variations, difficulties in aligning noncoding regions, and the presence of highly divergent sequences.

Over the past two decades, alignment-free approaches based on shared properties of sub-sequences of defined length *k* (*k*-mers or *k*-grams) (Marçais and Kingsford 2011; Shedlock et al. 2007; Chapus et al. 2005; Deschavanne et al. 1999) have been developed to compare sequences and genomes. Alignment-free methods for evolutionary analysis have been reviewed (Vinga and Almeida 2003; Zielezinski et al. 2019) and their robustness investigated (Chan et al. 2014; Bernard et al. 2016). For example, they can be used to derive distances to be summarized into phylogenies (Edwards et al. 2002; Chan et al. 2014; Balaban et al. 2022; Van Etten et al. 2023). Several studies have shown that alignment artifacts can significantly impact tree topology (Ogden and Rosenberg 2006; Wong et al. 2008; Du et al. 2019). Alignment becomes problematic with comparisons of large genomes, complex genomic variations, challenges in aligning noncoding regions, difficulties presented by highly divergent sequences, and the time-consuming nature of aligning large datasets. Alignment-free approaches offer a promising alternative to address these weaknesses of alignment-based methods (Ren et al. 2018).

Probabilistic topic modeling (Blei et al. 2003) is a statistical approach aiming to identify major ‘themes,’ ‘connections’, or ‘topics’ among themes in documents and other large collections of text. The approach originated from the field of Natural Language Processing (NLP) and was introduced by statisticians looking for applications of machine learning. Griffiths and Steyvers (2004) applied this method to infer a natural grouping (topics) of documents based on the content from a large number of scientific documents, called a corpus. Latent Dirichlet Allocation (LDA) is a popular technique in topic modeling introduced by Blei et al. (2003) within the context of unsupervised machine learning; it assumes that the documents consist of latent topics, each represented by the distribution of words. The goal is to uncover these topics by analyzing the words in the documents and essentially ‘learning’ the structure of the data. LDA has also been a focus of attention within the bioinformatics community and various applications to biological data have been researched and analyzed (Liu et al. 2016). Recently, some applications of alignment-free methods have been presented to solve problems offered by DNA sequences or genomes. For example, in a statistical application, LDA was used by La Rosa et al. (2015) to extract the frequency of fixed-length *k*-mers (words) of DNA sequences (documents) and thereby discover latent patterns in massive biological data to be used for clustering and classification of DNA sequences. Other studies have adopted LDA clustering for the analysis of single-cell expression or epigenetic data (duVerle et al. 2016; Dey et al. 2017). Here, we present a novel computational approach using probabilistic topic modeling to infer evolutionary relationships among individuals from different populations or species. This method works with multilocus data, including unaligned or aligned DNA sequences and unassembled raw sequencing reads. We will use the term species tree for these trees, whether derived from single individual sequences or individuals grouped into populations or species.

Genome-wide datasets (sequences of whole genomes or multiple genes per species) are becoming increasingly prominent in inferring the evolutionary history of closely related species. Traditional approaches to multilocus phylogenetics, such as concatenation methods (Gatesy and Baker 2005), and approaches that are consistent under the multispecies coalescent (de Queiroz and Gatesy 2007; Liu et al. 2009; Chifman and Kubatko 2014; Zhang et al. 2018; Mirarab et al. 2016), have advanced the field significantly. However, these methods often rely on high-quality alignments, which can be computationally expensive and error-prone when dealing with large or complex datasets.

TopicContml offers a scalable and efficient alternative to traditional approaches, addressing their limitations and enabling the analysis of diverse and complex genomic datasets without relying on sequence alignment.

Here, we outline the architecture of TopicContml and demonstrate its application using simulated data and three empirical datasets: (1) a small orthologous dataset of individuals from various locations of two parapatric bird species, consisting of 14 loci, along with a comparison to SVDquartets (Chifman and Kubatko 2014) and the alignment-free method Mash (Ondov et al. 2016), including a discussion of bootstrap support; (2) an orthologous dataset of mammalian species, consisting of 5,162 loci across 90 vertebrate species; and (3) a 12-species bird dataset with unaligned, non-orthologous PacBio raw sequencing reads.

## Materials and Methods

### TopicContml Software

TopicContml is a Python package based on a two-phase pipeline: (1) The mulitlocus or genome-wide data is broken down into *k*-mers; these *k*-mers are then used to learn a probabilistic topic model and extracts the topic frequencies of these *k*-mers using the LDA model for each locus (Fig. 1 left). For a data analysis with multiple individuals from the same species or population, we set the option `--merging n` to merge individual labels that start with the same *n* letters into groups; otherwise we assume each sequence is an individual. (2) These topic frequencies from multiple loci are then used to estimate a phylogeny with Contml (part of PHYLIP; Felsenstein 1981; 2004) (Fig. 1 right).

**FIGURE 1.**
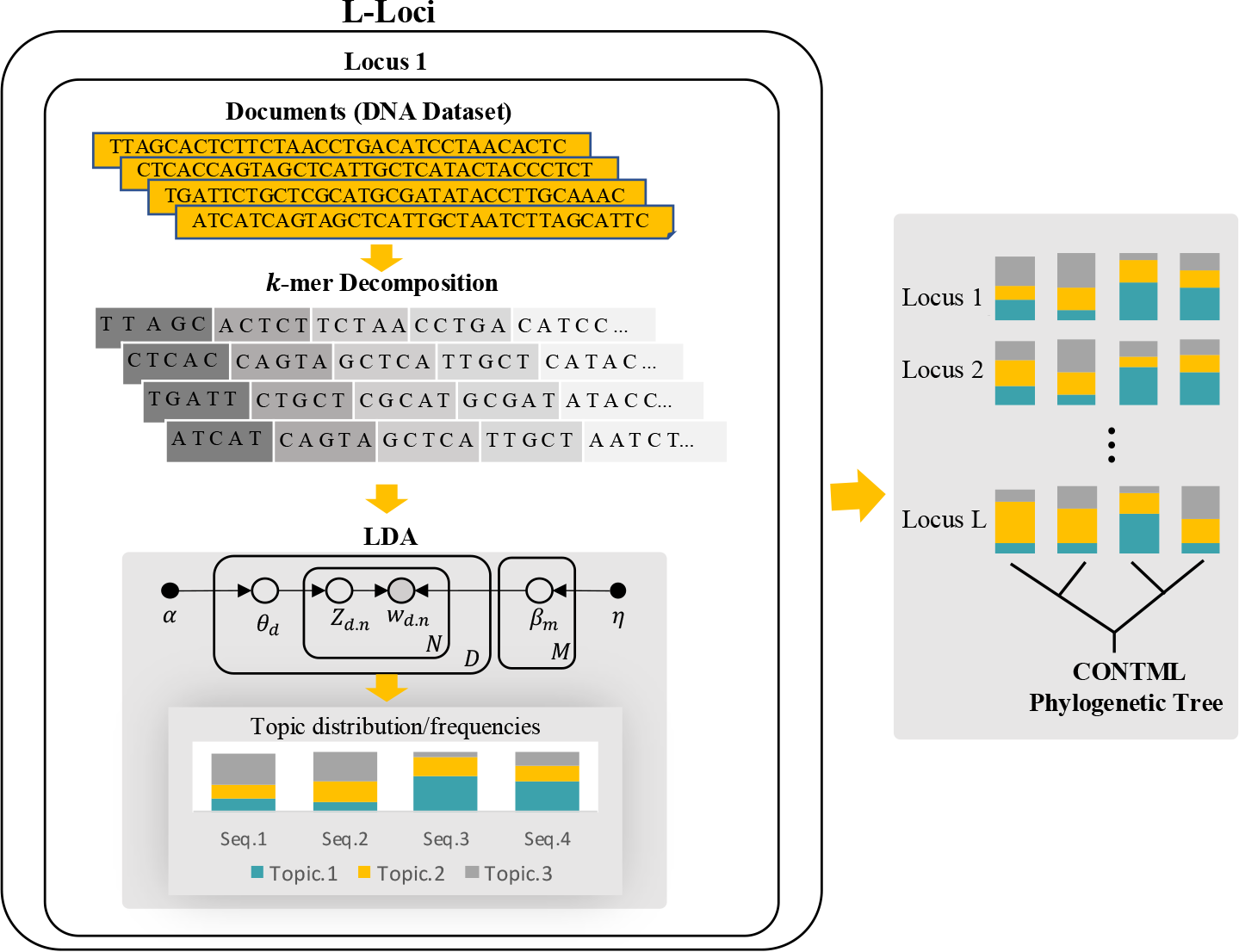
TopicContml workflow to generate topic frequencies and the corresponding phylogeny.

#### K-mer decomposition

In natural language processing (NLP), large text datasets (corpora) are broken down into smaller units such as documents, which are further divided into words or sentences, referred to as tokens. Similarly, in bioinformatics, datasets consisting of multiple genomes or multilocus DNA sequences can be broken down into individual genomes or groups of multilocus sequences associated with an individual. These sequences are then decomposed into *k*-mers—substrings of length *k* representing short DNA or amino acid sequences. We decompose the DNA sequences into non-overlapping *k*-mers, as shown in Fig. 1, since overlapping *k*-mers require more memory and computation time without yielding significantly better results. The program estimates an optimal *k*-mer length based on the probability of observing a given *k*-mer in a document (sequence) at each locus, with options for user adjustments.

The probability of a given *k*-mer *K* appearing in a random genome *X* of size *n* is *P* (*K ∈ X*) = 1 −(1 − |Σ|^−*k*^)^*n*^, where Σ = {*A, C, G, T*}, without loss of generality. Given a document size *n* and the desired probability *q* of observing a random *k*-mer, the value of *k* that minimizes the probability of observing a random *k*-mer can be computed as (Fofanov et al. 2004; Ondov et al. 2016):

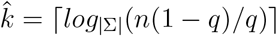

TopicContml calculates 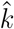 for each document in a locus based on this. We have found that *k* = 20 and *k* = 8 give accurate estimates in most cases for large (e.g., 1,000,000 bp) and small sequences, respectively. The program also allows users to choose different *k*-mer configurations, such as a combination of *k*-mer lengths or a single fixed length, based on their analysis needs.

#### Resolving ambiguities and missing data

The current version of TopicContml retains all IUPAC codes as they are, except for ‘N’, ‘?’, ‘-’, and other ambiguous characters, which can be filtered out prior to analysis. We assume that such ambiguity codes are rare and do not strongly affect the results.

#### Topic modeling

Given a collection of *D* documents and a number of *M* topics, topic modeling discovers the *M* topics from a collection of text data and estimates the probability of each topic for each document. We use Latent Dirichlet Allocation (LDA) to extract these frequencies. LDA is a generative probabilistic model used to uncover hidden topics within a collection of documents, referred to as a corpus. Given a corpus with *D* documents, let *N* be the number of words in a specific document *d ∈ D*. For each word *w*_*d,n*_ (the *n*^*th*^ word in the *d*^*th*^ document), *z*_*d,n*_ denotes the associated topic. The distribution of topics for document *d*, represented by *θ*_*d*_, is drawn from a Dirichlet distribution, *θ*_*d*_ *∼ Dir*(*α*), where *α* > 0 is the parameter vector. Similarly, the distribution of words for each topic *m*, denoted by *β*_*m*_, is also drawn from a Dirichlet distribution, *β*_*m*_ *∼ Dir*(*η*), with parameter vector *η* > 0. In this model, the only observed variables are the words *w* in the documents, while the topics *z*, the topic distributions *θ* for all documents, and the word distributions *β* for each topic are all latent variables. The joint distribution is defined as

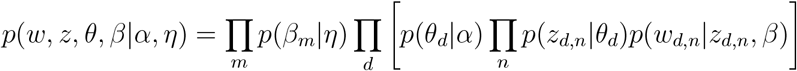

Using this joint distribution, one can compute the posterior distribution of the unknown model parameters, 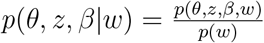 using expectation-propagation (EP) (Minka and Lafferty 2012) or other maximization methods.

For each locus, TopicContml first estimates the topic frequencies for every document, *θ*, using the Python package Gensim (Řehůřek and Sojka 2010) (Fig. 1). The sequences (documents) in each locus are decomposed into *k*-mers (words). During preprocessing, LDA filters out certain words, primarily those with low frequency, to enhance the quality of the generated topics (see Supplementary Fig. S8). The process begins by randomly assigning a distribution of topics to each document and a distribution of words to each topic, with these distributions being governed by Dirichlet priors. During the training phase, Gensim’s implementation of LDA iteratively refines these distributions using maximization methods. This training involves updating two key parameters: the distribution of topics within each document and the distribution of words within each topic. The model trains by analyzing patterns of word co-occurrence across the documents, assigning words to topics in a way that maximizes the likelihood of the observed data. As the iterations progress, the model converges to a stable set of topics.

#### Determining the optimal number of topics

Selecting the optimal number of topics in Latent Dirichlet Allocation (LDA) modeling is important for generating interpretable results. A widely used method for this involves evaluating topic coherence, which reflects the semantic similarity among top words within each topic—higher coherence scores generally correlate with more interpretable topics (Röder et al. 2015). Several algorithms are available for calculating coherence scores. In this study, we use the “*u*_*mass*_” coherence measure in Gensim (Řehůřek and Sojka 2010), analyzing each locus individually to identify the best topic number for each (see Supplementary Fig. S3).

However, because coherence analysis involves testing multiple models with varying topic counts, it can be time-intensive, especially with large datasets. To address this, TopicContml also allows users to specify a fixed number of topics, which, in our case, proved to be a practical alternative without compromising the interpretability or consistency of results. Although selecting an optimal topic count can enhance detail in some studies, our findings show that a fixed topic number performs well and offers efficiency for large-scale analyses. All our analyzed datasets are based on a fixed value of five topics.

#### Visualizing topics and associated terms

To visualize the topics and associated terms (*k*-mers), we use the package pyLDAvis (Sievert and Shirley 2014), an interactive visualization tool. TopicContml allows users to generate an HTML file containing the pyLDAvis output for each locus, saving them in a designated folder within the directory (see Supplementary Fig. S2). This enables detailed examination and extraction of information from the fitted LDA model, enhancing our ability to interpret the underlying topic structures

#### Contml (Continuous Characters Maximum Likelihood method)

The results of the LDA analysis are the topic frequencies for each document, which are then evaluated using Contml to estimate a phylogeny from frequency data using the restricted maximum likelihood method (REML) (Felsenstein 1973) based on the Brownian motion model for allele frequencies (Cavalli-Sforza and Edwards 1967). The primary assumption of Contml is that each character (each topic in our case) evolves independently according to a Brownian motion process and that character state changes since divergence are independent of each other, which means the net change after *t* units of time is normally distributed with zero mean and variance *ct* (the same constant *c* for all characters).

#### Multilocus bootstrapping

To assess the statistical confidence of the inferred phylogenies, we use bootstrapping (Efron 1979). Given a multiple sequence alignment, the bootstrap method involves resampling the original dataset with the replacement of the aligned sites and creating the phylogeny for each replicate. For unaligned data, we use the approach used in Edwards et al. (2002), where a random sample of *x k*-mers is drawn from all *x k*-mers collected from the data. The fraction of the time a particular clade appears in the resulting bootstrap trees presents support values for the clades in the reference tree or a majority-rule consensus tree (Holder et al. 2008; Efron et al. 1996; Hillis and Bull 1993). TopicContml implements bootstrapping strategies for both aligned and unaligned sequences. It generates a majority-rule consensus tree from the bootstrap replicates using SumTrees in DendroPy (Sukumaran and Holder 2010).

### Datasets

We evaluate the accuracy of TopicContml for simulated data and multiple real biological data. The simulated data were used to explore the effects of the number of loci and the accuracy of recovering the true topology from data sets consisting of 7 and 14 species. We used three biological datasets with very different features: (1) a 14-locus dataset from two parapatric closely related bird species separated into nine populations, to evaluate the accuracy of estimation, bootstrapping, and to compare accuracy with SVDquartets (Chifman and Kubatko 2014) and the alignment-free approach Mash (Ondov et al. 2016); (2) a vertebrate dataset focusing on mammals with 90 species and 5162 loci to evaluate the effect of missing data and of aligned vs. non-aligned orthologous loci; (3) a dataset of raw PacBio sequences of twelve bird species, each containing 100,000 reads; these sequences were neither orthologous nor aligned and can be construed as having been sampled randomly and potentially overlapping from their constituent genomes.

#### Simulated datasets

We evaluated two sets of simulations: one with moderate and one with many insertions and deletions. We used the software Dawg (Cartwright 2005) to simulate aligned sequences with indels/deletions on a 7-species and a 14-species tree (Fig. 2). The moderate scenario inserts indels at a rate of 0.02 per site and deletes sites with a rate of 0.02 per site. The more extreme simulation used an insertion/deletion rate of 0.2 per site. An example of two individual sequences for each indel/deletion scenario is shown in the electronic supplement (Supplementary Fig. S1). We simulated datasets of 1, 2, 5, 10, 20, 50, 100, 200, 500, and 1000 loci; each locus was between 800 - 2000bp, dependent on indels/deletions. These simulated datasets were then analyzed with gaps included (aligned), with the gaps excised (unaligned), and with *k*-mers that include gaps removed (no gap-kmer). We fixed the number of topics for all simulation analyses to 5 and estimated the best *k*-mer size from the data; it turned to be 9 for all datasets. Each scenario was run 10 times. The resulting trees were compared to the true trees using unweighted and weighted Robinson-Foulds distances.

**FIGURE 2.**
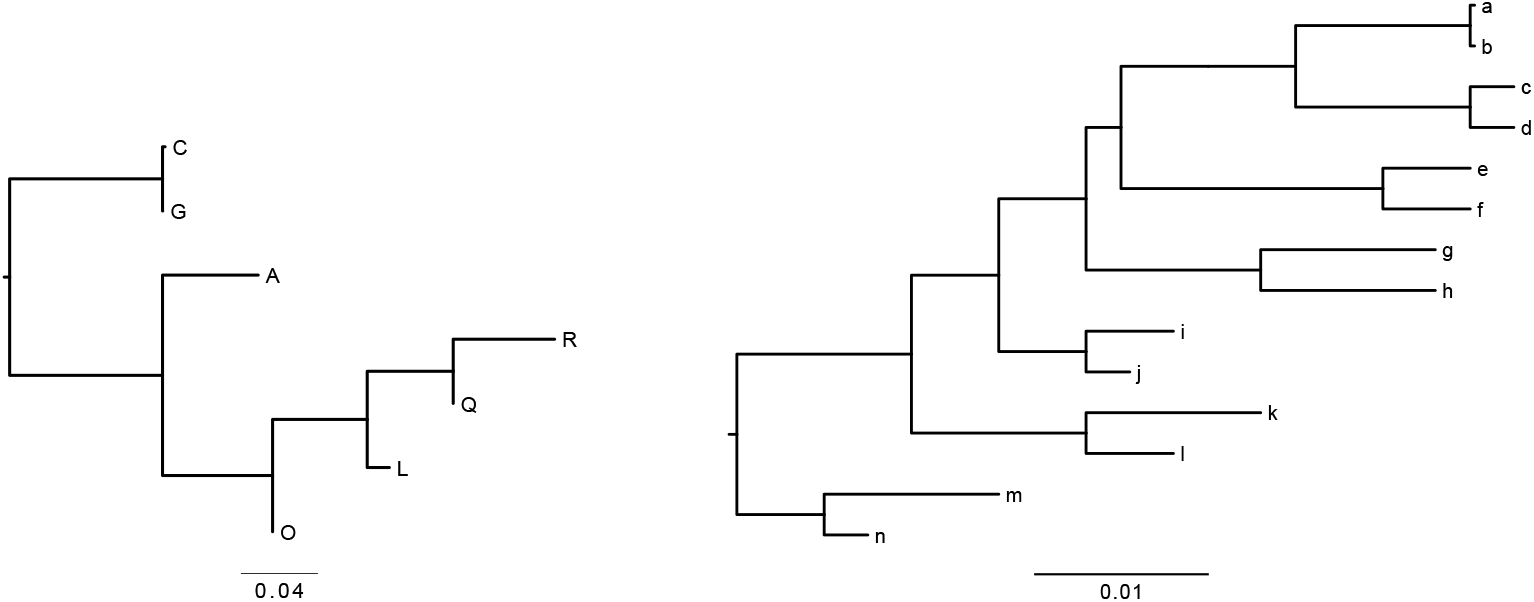
Phylogenies of seven and 14 species used for the simulation of sequence data with deletions and indels (gaps). In Tables 1 and 2, these are referenced as 7-tip and 14-tip trees.

#### Empirical datasets

We evaluated three biological datasets:

**Table 1.** Simulated data with low indel/deletion frequency, accuracy of phylogenetic reconstruction with TopicContml : data was simulated using Dawg (Cartwright 2005), each locus has around 1000 bp with gaps using a low number of gap scenario (parameter: insertion/deletion probability 0.02/site, average indel length 12, topics=5, *k*-mers were estimated from the data (all estimated *k*-mers were 9 base pairs); ‘Aligned’ results are based on simulated data with gaps, ‘Not Aligned’ had all gaps removed in the simulated data, ‘No gap-kmer’ had all *k*-mers containing gaps removed before LDA. ‘C’ (Close) marks the frequency of topologies that are either the same as the true topology or not more than 2 rearrangments apart; ‘wRF’ is the average weighted Robinson-Foulds distance from the true tree. Each result is based on 10 simulations.

| Loci | 7 Species |  |  |  |  |  | 14 Species |  |  |  |  |  |
| --- | --- | --- | --- | --- | --- | --- | --- | --- | --- | --- | --- | --- |
|  | Aligned |  | Not Aligned |  | No gap-kmer |  | Aligned |  | Not Aligned |  | No gap-kmer |  |
|  | C | wRF | C | wRF | C | wRF | C | wRF | C | wRF | C | wRF |
| 1 | 0.1 | 1.22 | 0.3 | 1.14 | 0.3 | 1.23 | 0.0 | 1.76 | 0.0 | 1.78 | 0.0 | 1.81 |
| 2 | 0.4 | 1.30 | 0.4 | 1.23 | 0.5 | 1.22 | 0.0 | 1.97 | 0.0 | 2.14 | 0.0 | 1.95 |
| 5 | 0.6 | 1.19 | 0.3 | 1.27 | 0.5 | 1.26 | 0.0 | 1.93 | 0.1 | 2.16 | 0.2 | 1.99 |
| 10 | 1.0 | 1.18 | 0.6 | 1.21 | 0.5 | 1.19 | 0.4 | 2.01 | 0.1 | 2.13 | 0.4 | 1.93 |
| 20 | 0.7 | 1.18 | 0.5 | 1.16 | 0.6 | 1.18 | 0.6 | 2.01 | 0.8 | 2.18 | 0.7 | 1.99 |
| 50 | 0.9 | 1.15 | 0.8 | 1.18 | 0.8 | 1.17 | 0.8 | 1.99 | 0.7 | 2.15 | 1.0 | 1.99 |
| 100 | 1.0 | 1.16 | 0.7 | 1.23 | 0.9 | 1.17 | 0.9 | 1.99 | 1.0 | 2.11 | 0.9 | 1.99 |
| 200 | 1.0 | 1.18 | 0.8 | 1.22 | 0.9 | 1.17 | 0.9 | 2.01 | 1.0 | 2.12 | 0.8 | 1.98 |
| 500 | 1.0 | 1.17 | 1.0 | 1.18 | 1.0 | 1.17 | 1.0 | 1.97 | 1.0 | 2.14 | 1.0 | 1.97 |
| 1000 | 1.0 | 1.17 | 0.9 | 1.22 | 1.0 | 1.18 | 1.0 | 1.96 | 1.0 | 2.13 | 1.0 | 1.95 |

**Table 2.** Simulated data with high indel/deletion frequency, accuracy of phylogenetic reconstruction with TopicContml : data was simulated using Dawg (Cartwright 2005), each locus has around 1000 bp with gaps using a high number of gap scenario (parameter: insertion/deletion probability 0.2/site, average indel length 12, topics=5; *k*-mers were estimated from the data (all estimated *k*-mers were 9 base pairs); ‘Aligned’ results are based on simulated data with gaps, ‘Not Aligned’ had all gaps removed in the simulated data, ‘No gap-kmer’ had all *k*-mers containing gaps removed before LDA. ‘C’ (Close) marks the frequency of topologies that are either the same as the true topology or not more than 2 rearrangments apart; ‘wRF’ is the average weighted Robinson-Foulds distance from the true tree. Each result is based on 10 simulations.

| Loci | 7 Species |  |  |  |  |  | 14 Species |  |  |  |  |  |
| --- | --- | --- | --- | --- | --- | --- | --- | --- | --- | --- | --- | --- |
|  | Aligned |  | Not Aligned |  | No gap-kmer |  | Aligned |  | Not Aligned |  | No gap-kmer |  |
|  | C | wRF | C | wRF | C | wRF | C | wRF | C | wRF | C | wRF |
| 1 | 0.5 | 1.33 | 0.1 | 1.24 | 0.1 | 1.27 | 0.0 | 1.91 | 0.0 | 2.08 | 0.0 | 1.59 |
| 2 | 0.2 | 1.17 | 0.2 | 1.26 | 0.5 | 1.27 | 0.0 | 2.08 | 0.0 | 3.08 | 0.0 | 2.29 |
| 5 | 0.6 | 1.29 | 0.2 | 1.32 | 0.6 | 1.18 | 0.3 | 2.04 | 0.0 | 3.36 | 0.2 | 2.05 |
| 10 | 0.6 | 1.22 | 0.4 | 1.18 | 0.4 | 1.19 | 0.2 | 2.05 | 0.0 | 3.66 | 0.4 | 2.04 |
| 20 | 0.8 | 1.20 | 0.7 | 1.21 | 0.6 | 1.24 | 0.7 | 2.11 | 0.0 | 3.78 | 0.5 | 2.06 |
| 50 | 0.7 | 1.21 | 0.9 | 1.19 | 0.5 | 1.23 | 0.9 | 2.09 | 0.0 | 3.74 | 0.7 | 2.03 |
| 100 | 0.5 | 1.25 | 0.8 | 1.22 | 0.5 | 1.22 | 1.0 | 2.09 | 0.0 | 3.74 | 0.8 | 2.01 |
| 200 | 0.7 | 1.22 | 0.8 | 1.23 | 0.6 | 1.23 | 1.0 | 2.08 | 0.0 | 3.81 | 0.9 | 1.99 |
| 500 | 0.9 | 1.20 | 1.0 | 1.22 | 0.3 | 1.24 | 1.0 | 2.06 | 0.0 | 3.85 | 1.0 | 1.99 |
| 1000 | 0.7 | 1.22 | 1.0 | 1.22 | 0.7 | 1.21 | 1.0 | 2.06 | 0.0 | 3.91 | 1.0 | 1.98 |

##### Two parapatric closely-related bird species

The data contain multiple individuals of the Australian brown treecreeper (*Climacteris picumnus*) and black-tailed treecreeper (*Climacteris melanurus*) (Edwards et al. 2022; 2023). DNA sequences consisted of 14 loci and nine different geographic locations. For each locus, sequence length varied from 288 to 418 base pairs, and the number of aligned sequences per locus ranged from 78 to 92. For the evaluations of unaligned data, we removed all gaps in each sequence. We applied LDA to each locus across all 9 locations and extracted the topic frequencies. These topic frequencies were used in Contml to generate the population tree, evaluate bootstrap support, and compare with another approach.

##### Mammal dataset

The second dataset includes 90 vertebrate species focusing on mammals with 5162 loci (Wu et al. 2018), and were analyzed by Liu et al. (2017) using concatenated maximum likelihood and coalescent approaches. We analyzed this dataset using TopicContml under four conditions: (a) excluding *k*-mers containing gaps (‘-’) or unspecified nucleotides (‘N’), (b) removing alignment columns with gaps before excluding *k*-mers with ‘N’, (c) removing all gaps from sequences before excluding *k*-mers with ‘N’, and (d) using aligned sequences while retaining ‘N’ characters. We then used the Robinson-Foulds distance to compare each phylogram with the maximum-likelihood tree and random trees.

##### PacBio dataset

A total of 6.5 GB of raw sequencing reads of 12 bird species generated by the PacBio HiFi sequencing method. The 12 birds cover most of the depth of the avian tree, including the two deepest branches, Paleognathae and Neognathae Species were chosen so as to broadly sample the tree for species belonging to lineages whose higher relationships are fairly stable, but also to include unambiguously close relatives, so that we could test the ability of our method to recover close relatives. We subsampled 100,000 reads from each species; average read length per species varied from 9.3 kb in the tinamou Crypterellus tataupa to 18.8 kb in chicken.

## Results

### Analysis of Simulated Datasets

The simulations based on the moderate insertion/deletion scheme, shown in Table 1, demonstrate that the recovery of trees close to the true tree improves with the number of loci for all treatments and also for both tree topologies. The ‘Aligned’ treatment for both tree topologies works well with more than 50 loci. The ‘Not aligned’ treatment worked better for the 14-species tree than for the 7 species tree, but we used only a small number of replicates (n=10). Still, accurate recovery of the true tree was almost as high as with the ‘Aligned’ treatment. The ‘No gap-kmer’ treatment fairs as well as the ‘Not aligned’ treatment. Weighted Robinson-Foulds distances (wRF) show trends consistent with the percentage of trees recovered that either match the true tree or are within two distance units of it. Notably, the ‘No gap-kmer’ treatment is closer to the true tree than the ‘Not aligned’ treatment, which performs comparably to the ‘Aligned’ treatment.

The simulations with extreme insertion/deletion strategy (as shown in Table 2) are markedly different for the ‘Not Aligned’ treatment of the 14 tip trees: even with 1000 loci none of the estimated trees were close to the true tree; ‘Aligned’ and ‘No gap-kmer’ faired similar to the moderate indel/deletion scheme. For the 7-species tree all treatments are close to the true tree when the number of loci is large. Overall the wRF values are all higher than those for the moderate indel/deletion scenario, and the ‘No gap-kmer’ delivers similar values as the ‘Aligned’.

### Analysis of Empirical Datasets

#### Multilocus species tree from closely related Australian birds

For the treecreeper dataset, we tokenized each DNA sequence at every locus using *k*-mer representation with a *k* value of 8 (as estimated by TopicContml), employing non-overlapping tokens. Individuals from the same location were merged, and LDA was applied to the corpus to generate topic frequencies for each locus for each of the 9 populations. We used five topics in our analysis. These multilocus topic frequencies were then used to construct a maximum-likelihood tree with Contml in the PHYLIP package. We did bootstrapping, and Fig. 3b shows the majority-rule consensus tree generated by TopicContml for unaligned data, with bootstrap support values derived from 1000 replicates. Inspection of the clades reveals that our tree recover the expected geographic relationship within each species (Edwards et al. 2023; Cracraft 1986), and the locations are separated by species, which in turn are separated by the Carpentarian barrier in Australia.

**FIGURE 3.**
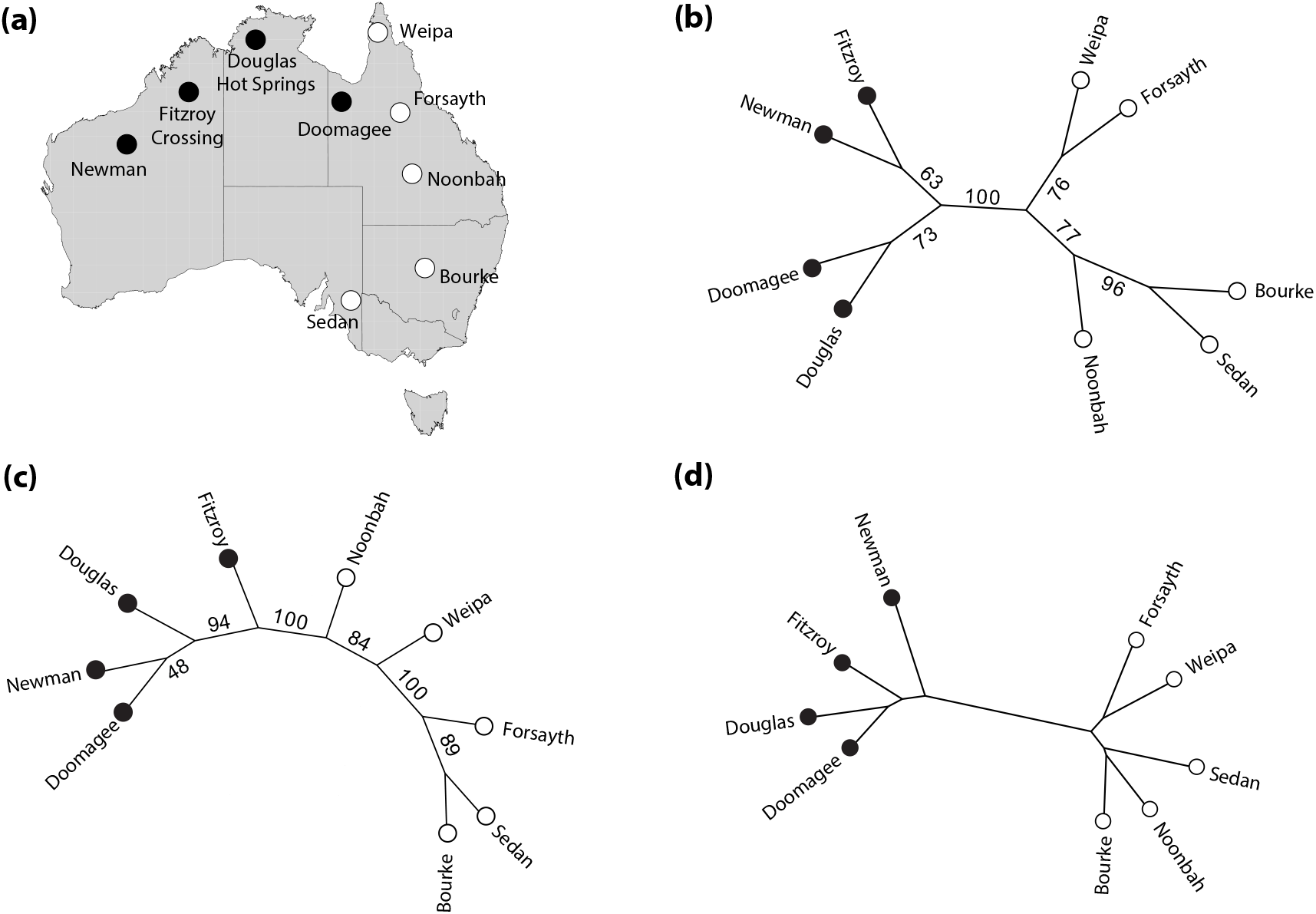
Relationship tree of 9 populations of two Australian treecreeper species reconstructed. (a) The map of Australia shows the locations; the black disks mark *Climacteris melanurus*, and the white disks mark *Climacteris picumnus* (Edwards et al. 2023). (b) The majority-rule consensus tree of unaligned data by TopicContml. For each bootstrap analysis, 1000 replicates were used. Values in the graphs are % support. (c) The majority-consensus tree of aligned data analyzed by SVDquartets +Paup*. (d) The phylogeny constructed by Mash.

We compared the performance of TopicContml with SVDquartets (Chifman and Kubatko 2014) implemented in Paup* (Swofford 2003) (SVDquartets +Paup*), as shown in Fig. 3c. There are notable differences between the results of SVDquartets and TopicContml When comparing these results to the mapped locations in Fig. 3a, we observe that our bootstrap tree from TopicContml recovers the relationships equally well or better than SVDquartets. Both methods encounter challenges in resolving certain population splits but confidently separate the two species. TopicContml support values recover, in general, the geographic pattern of the locations well, and a greater number of potential splits are unresolved.

We also compared our phylogenetic tree with one generated using the alignment-free approach Mash (Ondov et al. 2016), which estimates evolutionary distances between nucleotide sequences. For the Mash input, sequences from different loci for each individual were concatenated, with missing data filled by gaps. The sequences from individuals at the same location were then merged to create nine Fasta files, representing the nine populations in Australia. The pairwise distance matrix generated by Mash was used to construct a Neighbor-Joining tree using the PHYLIP package (Felsenstein 2004). Fig. 3d shows the phylogenetic tree generated by Mash. When comparing our tree to the one generated by Mash, we observe that both trees depict similar relationships among species.

#### Multilocus species tree from genome-wide mammal dataset

We analyzed a mammal data with 90 species and 5162 loci. The dataset consisted of nucleotide characters from the set ‘ACGT-N’. We analyzed the mammal dataset under various conditions and compared the resulting trees to the maximum likelihood tree derived from 4388 loci of 90 vertebrate species, as reported by Liu et al. (2017). The comparisons were visualized using tanglegrams (Revell 2024). In the first analysis, after generating *k*-mers, we excluded any *k*-mers containing either ‘-’ or ‘N’. The resulting phylogenetic tree (Fig. 4) was 60 steps away from the maximum likelihood tree, as measured by the Robinson-Foulds (RF) distance. In the second analysis, we removed all alignment columns containing gaps, then excluded *k*-mers with ‘N’. The resulting tree (Supplementary Fig. S4) was 64 steps from the reference tree. This resulted in a tree generated using 1719 loci because 3443 of the 5162 loci did not contain any data after the removal. In the third analysis, we removed all gaps from each sequence before excluding *k*-mers containing ‘N’. This approach did not improve the tree (Supplementary Fig. S5), which was 90 steps from the reference tree. Finally, in the fourth analysis, we generated a tree using aligned sequences while retaining ‘N’ characters. This tree (Supplementary Fig. S6) was 80 steps from the maximum likelihood tree.

**FIGURE 4.**
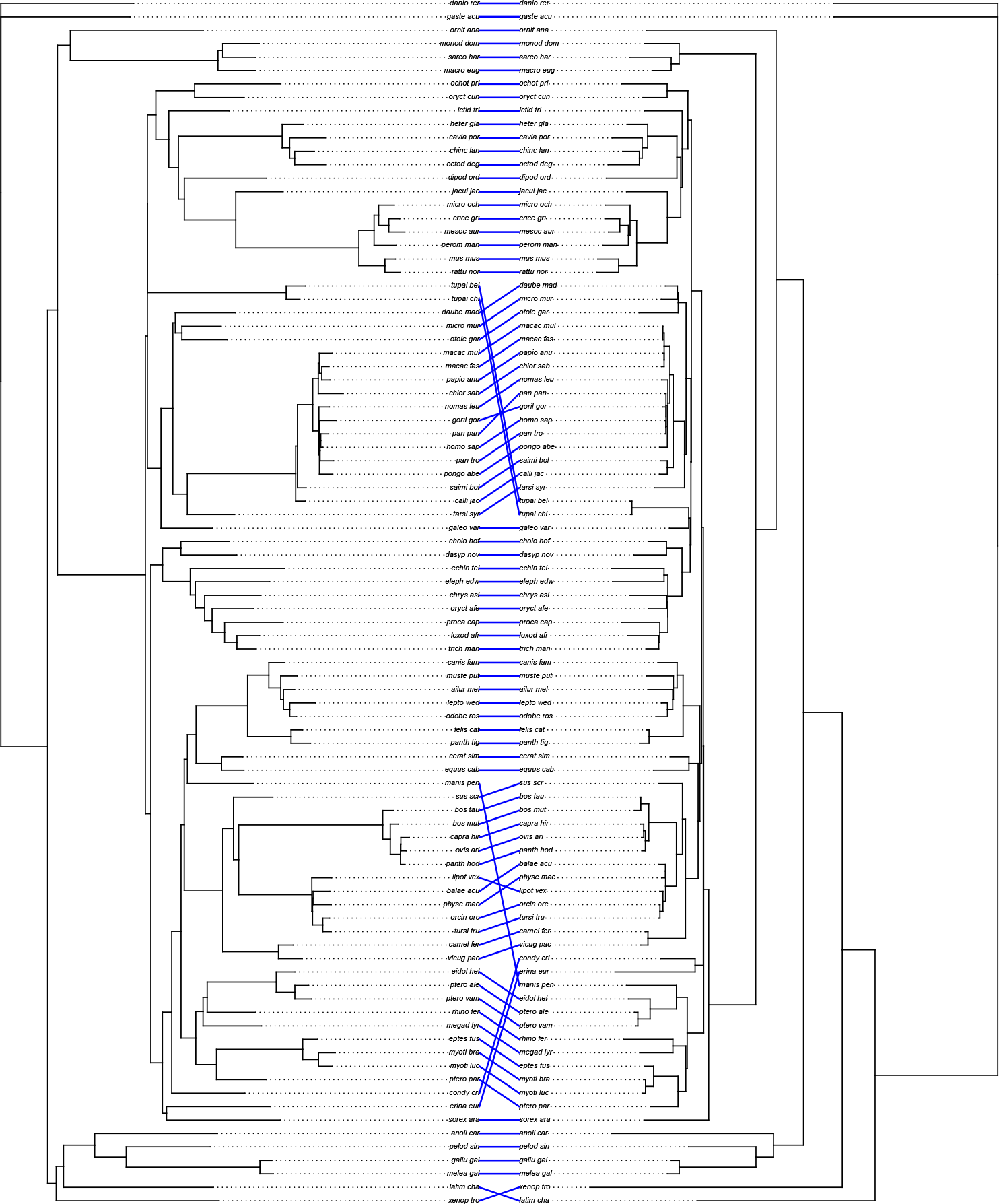
Tanglegram of mammal dataset comparing the TopicContml tree (left), generated by excluding *k*-mers containing ‘-’ or ‘N’, with the maximum likelihood tree from Liu et al. (2017) (right). The alphabetical list of the species names in the tree is in Supplementary Table S1.

The RF distances between 1,000,000 random trees and the maximum likelihood tree reveal a minimum distance of 168, a mean distance of 173.5, and a maximal distance of 174. The topic modeling trees are therefore considerably closer to the maximum likelihood tree than random trees (Supplementary Fig. S7), confirming that our topic modeling approach recovers phylogenetic signal. The tanglegrams (Fig. 4; Supplementary Fig. S4, S5, S6) also confirm that our tree and the maximum likelihood tree are fairly similar despite a seemingly large RF distance, especially when considering that many of the branches in the mammal tree are very short.

#### Multilocus species tree from raw unassembled PacBio sequence reads

This dataset consists of FASTA files containing 100,000 reads from each of 12 species. To reduce computational time by reducing the number of loci, we first combined sets of 1,000 reads and then concatenated them to produce files of 100 loci. Fig. 5 (left) displays the tree generated by TopicContml using a non-overlapping *k*-mer length of 20, with an RF distance of two from the reference tree, as shown in Fig. 5 (right). The reference tree was drawn from Cracraft (1988), Oliveros et al. (2019), and Wu et al. (2024). The discrepancy in relationships of the Wrentit and Yellow warbler are uncertain, because relationships in this portion of the passerine tree are certainly not definitive.

**FIGURE 5.**
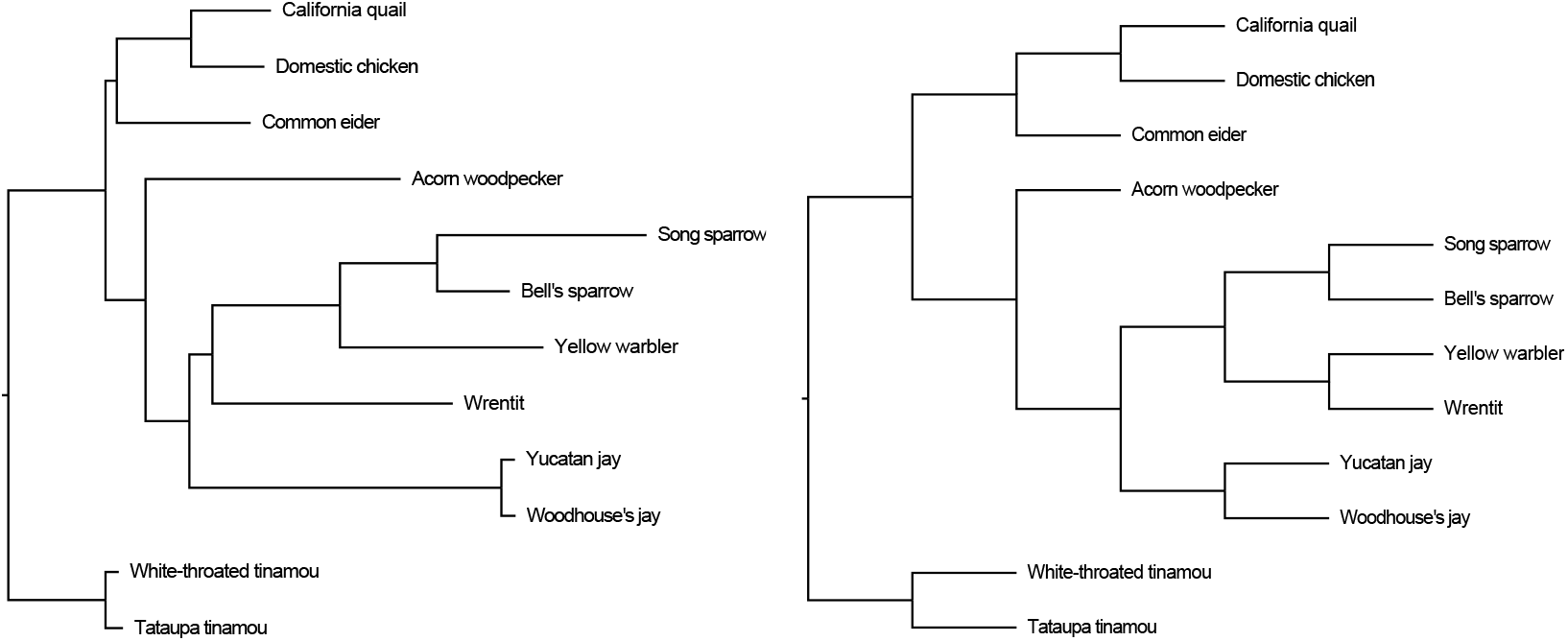
The phylogeny generated by TopicContml (left side) compared to the reference tree (right side).

Given the large size of each document, we performed our first analysis using a longer *k*-mer length (*k*-mer length of 20 as we discussed in Section *K*-mer decomposition). To assess the effects of different *k*-mer ranges, we experimented with lengths from 8 to 42. We found that optimal results were achieved with *k*-mers in the range of 18 to 30, as demonstrated in Supplementary Fig. S9.

## Discussion

This study integrates *k*-mers and probabilistic topic modeling to perform phylogenetic analysis on unaligned or aligned multilocus sequence data, as well as on unassembled raw sequencing reads. The Python code TopicContml offers an efficient workflow to reconstruct evolutionary relationships potentially without prior sequence alignment. TopicContml contains a two-phase analysis. First, *k*-mers are extracted from DNA sequences, and Latent Dirichlet Allocation uses these *k*-mers to establish how probable an individual’s set of *k*-mers fits an arbitrary number of topics. For each locus and each individual, we generate a vector of assignment frequencies for a predefined set of topics. This step can be parallelized among loci. The LDA runtime depends on the length of the sequences (documents) and the number of loci. In the second step, the topic frequencies are used as input for Contml to construct a phylogenetic tree. The Contml evaluation time is influenced by the number of tips and the number of loci, as the input matrix for Contml is structured as number of tips × (number of loci × (number of topics - 1)). Although the current version of Contml does not support parallelization, implementing this capability could significantly improve its runtime, particularly for analyses involving many species. As shown in Supplementary Table S3, the Australian bird dataset completed in seconds for both LDA and Contml. The mammal dataset, however, took longer for LDA due to its large number of loci (5,162), despite relatively short sequence lengths, and even longer for Contml due to the large input size. For the PacBio dataset, the LDA runtime was extended by the significantly longer sequence lengths, but the Contml step completed within seconds.

The simulated data experiments reflect the influence of gap handling on phylogenetic reconstruction accuracy under different insertion/deletion (indel) rates and tree complexities. The simulation protocol produced aligned data with low and high gap numbers. The more aligned the data was, the more accurate was the phylogoenetic inference; for both moderate and high indel frequencies, this approach consistently outperforms others. The observed decline in accuracy for the ‘Not Aligned’ group, especially in the high-indel 14-species tree, stems from the loss of indel information because gaps are removed entirely; new *k*-mer were then formed that do not necessarily coincide with the phylogenetic signal. The ‘Not Aligned’ group faired well with the moderate indel scheme because only few *k*-mers were affected. The ‘No gap-kmer’ group’s results, which approximate those of the ‘Aligned’ group under moderate indel scenarios, suggest that excluding only gap-containing *k*-mers strikes a balance between noise reduction and information preservation. This selective approach retains enough informative content while minimizing the alignment artifacts that become problematic in high-indel scenarios.

In contrast, the analysis of the real datasets was more complex, as demonstrated using the mammal dataset to evaluate the effect of structuring sequence data into *k*-mers. The Robinson-Foulds (RF) distances to the maximum likelihood tree (Liu et al. 2017) reveal that increasing the number of loci does not always improve phylogenetic accuracy. The mammal dataset analysis highlights the robustness of TopicContml in recovering phylogenetic signal under various treatments of gaps and missing data. The most accurate tree (RF distance of 60) was achieved by excluding *k*-mers containing gaps or ‘N’, underscoring the importance of targeted ambiguity removal. Removing entire alignment columns with gaps reduced the number of loci but still produced a comparable tree (RF distance of 64). Conversely, removing all gaps from sequences resulted in the least accurate tree (RF distance of 90), likely due to the loss of biologically informative gap signals. Retaining ‘N’ characters in aligned sequences yielded an intermediate result (RF distance of 80), showing that while ambiguity introduces noise, key phylogenetic relationships are still preserved. Notably, all TopicContml trees were substantially closer to the reference tree than random trees.

Tree uncertainty is commonly assessed through bootstrap analysis, which poses challenges for unaligned datasets as it requires bootstrapping at the *k*-mer level rather than the sequence level. For the treecreeper dataset, bootstrap analysis with TopicContml demonstrates its robustness in recovering phylogenetic relationships from unaligned data. The majority-rule consensus tree effectively separates the two species and accurately captures geographic patterns, including the division across the Carpentarian barrier. Compared to the alignment-based SVDquartets, TopicContml achieves equal or better precision in recovering geographic relationships. Furthermore, the comparison with the alignment-free method Mash confirms that TopicContmleffectively captures key phylogenetic relationships while working with unaligned data.

Our analyses of the PacBio dataset shows substantial promise deriving phylogenetic signal from unaligned long-read sequences, and demonstrates the potential of TopicContmlfor alignment-free phylogenetic reconstruction. Despite the complexity of raw, unassembled reads, TopicContml produced trees closely matching a reference phylogeny, showing its capacity to infer evolutionary relationships directly from complex, heterogeneous data. An important goal of the future is to determine what components of unaligned genomic data - tranposable elements, satellite sequences, or other common components of genomes - are driving these positive results. Although the LDA step for this data set required additional processing time due to long-read lengths, Contml efficiently completed tree inference. These results suggest that TopicContml offers a promising approach for handling high-throughput phylogenetic data without requiring sequence alignment or even genome or locus assembly.

TopicContml is modular, and we have begunwork to replace Contml with a network-generating package that may improve the analyses of such datasets by incorporating gene flow between species. Currently, for many datasets that do not suffer widespread introgression, TopicContml allows the analysis of many loci from many individuals that can be grouped, for example, into locations or species. We believe that TopicContml will become a valuable addition to the computational toolkit for phylogenetics by constructing evolutionary trees without or with sequence alignment.

## Software Availability

We implemented our method as a free software named TopicContml under the MIT open-source license. The source code and the documentation of TopicContmlare available at https://github.com/TaraKhodaei/TopicContml.git

## Acknowledgments

We sincerely thank the reviewers for their constructive comments and suggestions, which significantly improved this manuscript. We acknowledge the contributions of our collaborators and are grateful to Subir Shakya and Tim Sackton for providing pre-publication access to PacBio reads from the two tinamou species. Additionally, we thank Liang Liu for assistance with interpreting the mammal dataset. We also thank the Louisiana State University Museum of Science for access to the *Tinamus guttatus* specimen used to generate PacBio sequence data, and the staff of the Museum of Comparative Zoology Ornithology Department for access to the jay specimens. Some simulations were conducted on the Research Computing Cluster at Florida State University, Tallahassee, and some data preprocessing was performed on the FASRC Cannon cluster, supported by the FAS Division of Science Research Computing Group at Harvard University.

## Data Availability

- **Simulated Data**: Instructions for generating the simulated data are available at https://github.com/pbeerli/simulations_for_topiccontml.
- **Australian Treecreeper Dataset**: Available through Dryad (Edwards et al. 2022).
- **Mammal Dataset**: The aligned mammal dataset can be accessed at https://figshare.com/articles/cds_5162_zip/6031190 (Wu et al. 2018).
- **PacBio Bird Sequences**: The sources of PacBio long-read sequences from birds are detailed in Supplementary Section: Processing of PacBio Reads and Supplementary Table S2. Datasets for the four unpublished species (two tinamous and two jays) are available through Dryad at https://datadryad.org/stash/share/5RpvXGzGfUdffDrC3Jv_LF8oSvuY3xiIufQDgn5YELg.

## Electronic Supplement

### Simulated dataset

We evaluated two sets of simulations: one with moderate and one with many insertions and deletions. We used the software Dawg (Cartwright 2005) to simulate aligned sequences with indels/deletions on a 7-species and a 14-species tree (Fig. 2). The moderate scenario inserts indels at a rate of 0.02 per site and deletes sites with a rate of 0.02 per site; for example, in the 7-species simulations, a species could have at a locus on average 1038 sites that include, on average per species 8.2 gaps with and average length of 12 sites. The more extreme simulation used an insertion/deletion rate of 0.2 per site. The number of gap sites increased for the specific tree considerably: each locus had about 59.3 gaps on average 17 sites long, each species had about 960 A, C, G, T sites, and about 1000 ‘-’ sites. The branch length and topology matter for the gap distribution: the 14-tip tree had a similar distribution of the gaps with the low insertion/deletion parameter as the 7-tip tree, but the high parameter resulted in many gaps but not as extreme as in the 7-tip tree (about 400 ‘-’ sites per species).

**Figure S1.**
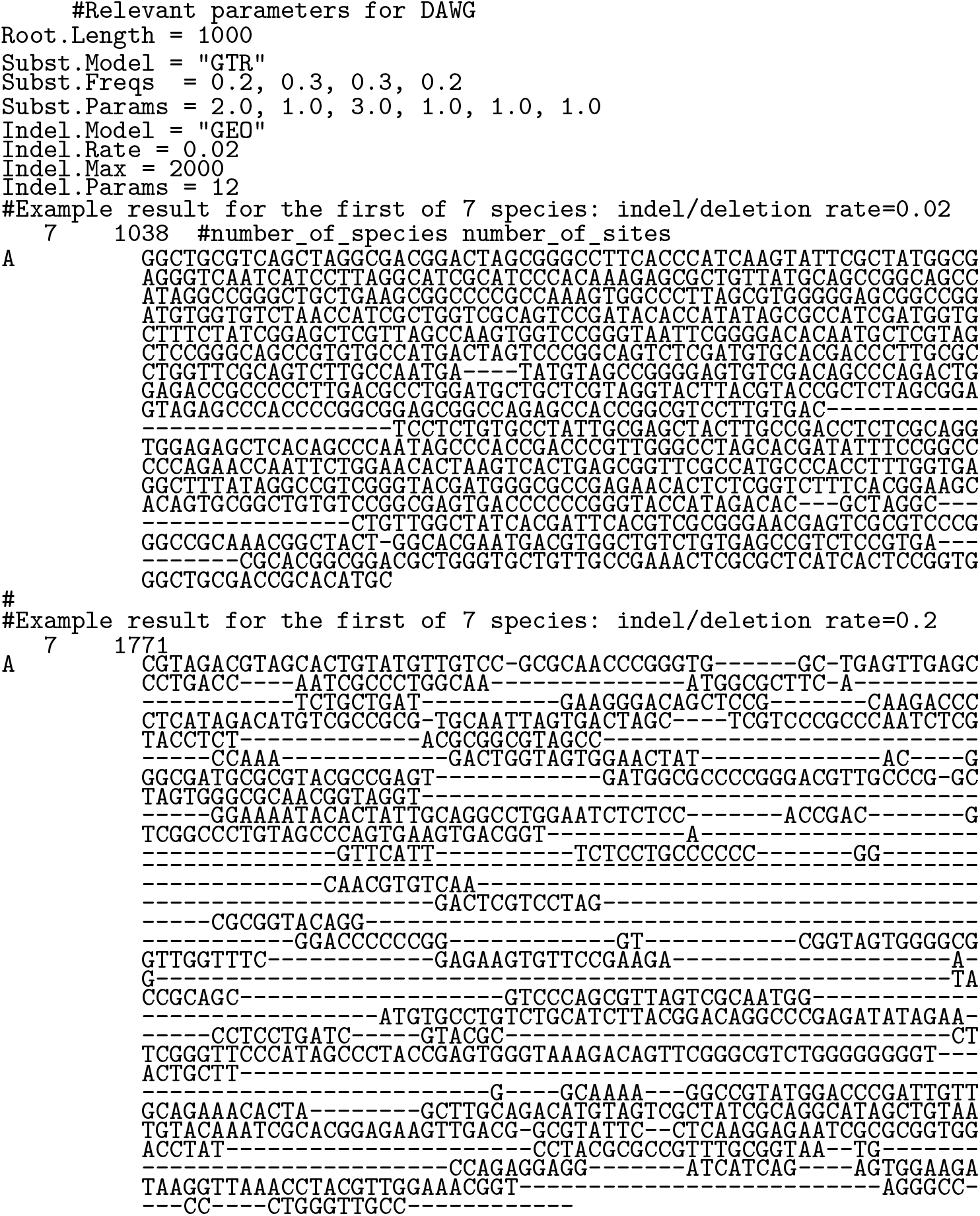
Example sequences for the two simulation treatement, resulting in data that contain gaps determined by the deletion/indel parameter in the simulation software DAWG Cartwright (2005).

### Bird dataset

**Figure S2.**
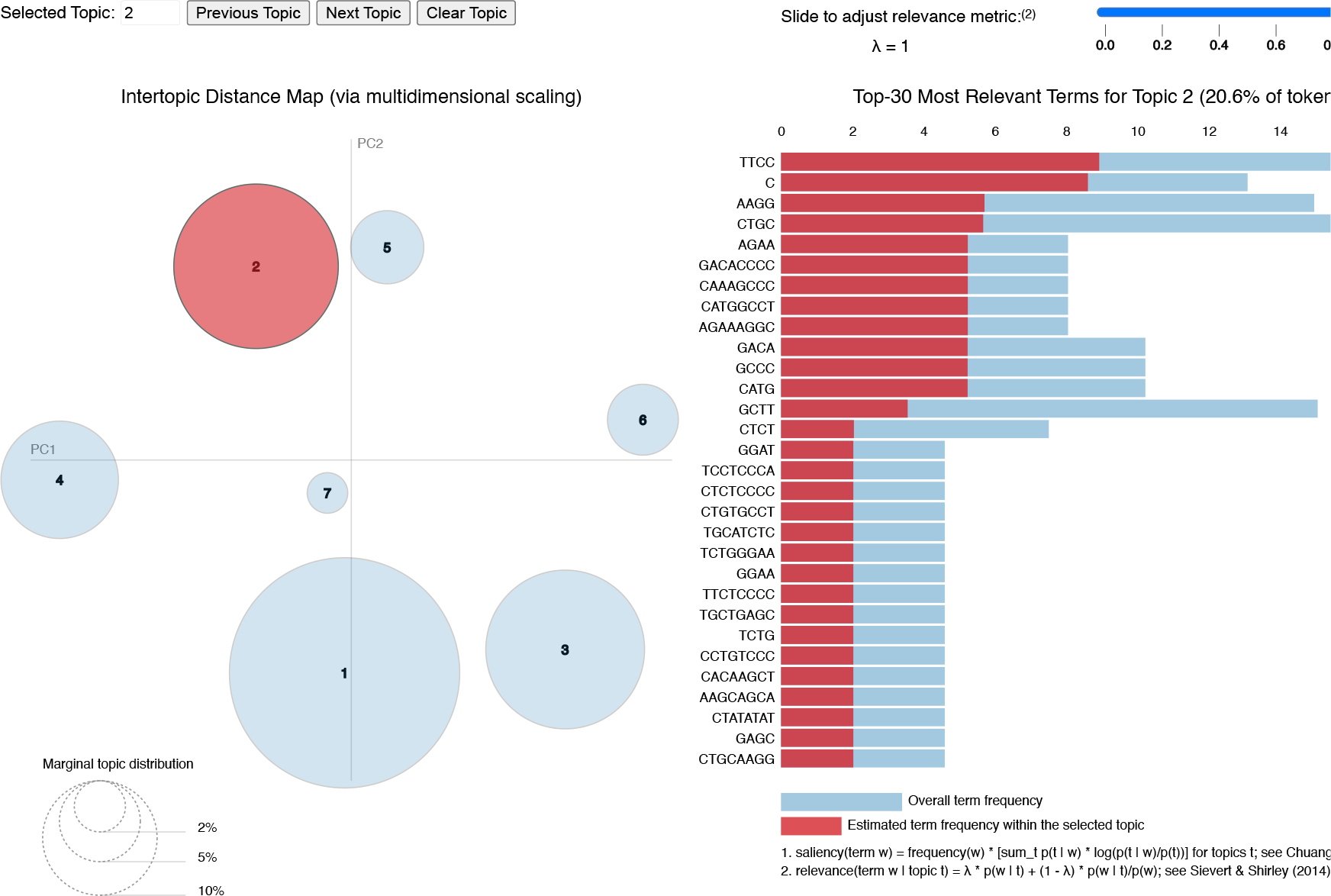
Screenshot of an interactive two-part visualization output from pyLDAvis for the ‘Bird’ dataset, first locus. Left: an intertopic distance map of topics (7 topics) generated by the LDA model for the first locus. The topics are represented as circles scaled to their frequency of occurrence. The index number of each cluster represents the topic ID; the red circle shows the current topic, and the blue circles show other topics. Right: distribution of the top 30 most relevant terms among topics.

**Figure S3.**
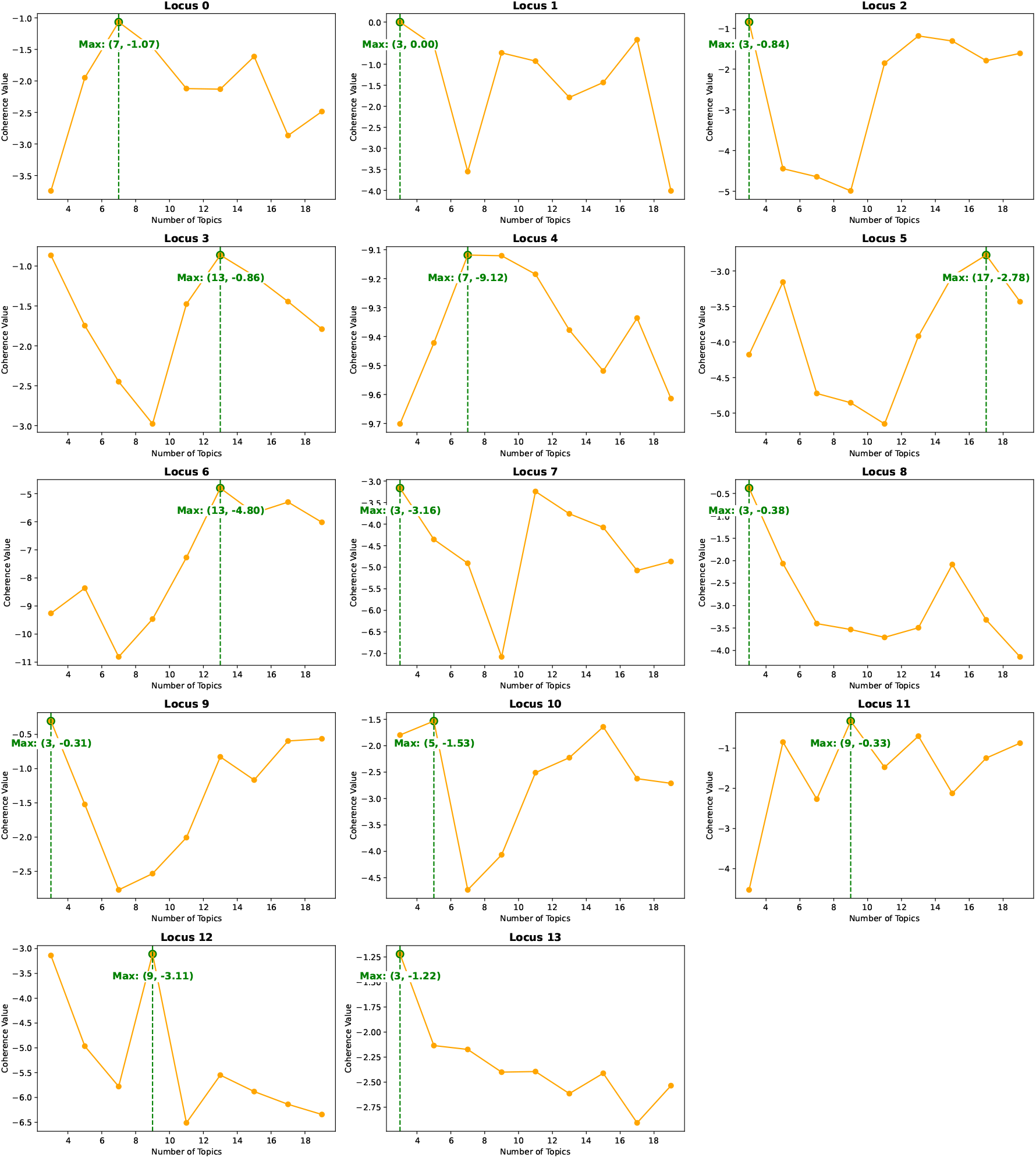
Coherence scores for varying numbers of topics in the range of 3 to 20 (with an increment of 2) for each of the 14 loci in the ‘Bird’ dataset. Each subplot represents the coherence analysis for a specific locus, illustrating the *u mass* coherence scores for topic numbers 3, 5, 7, 9, 11, 13, 15, 17, and 19. Higher coherence scores indicate more interpretable topics, allowing us to determine the optimal number of topics for each locus individually.

### Mammal dataset

**Table S1.**
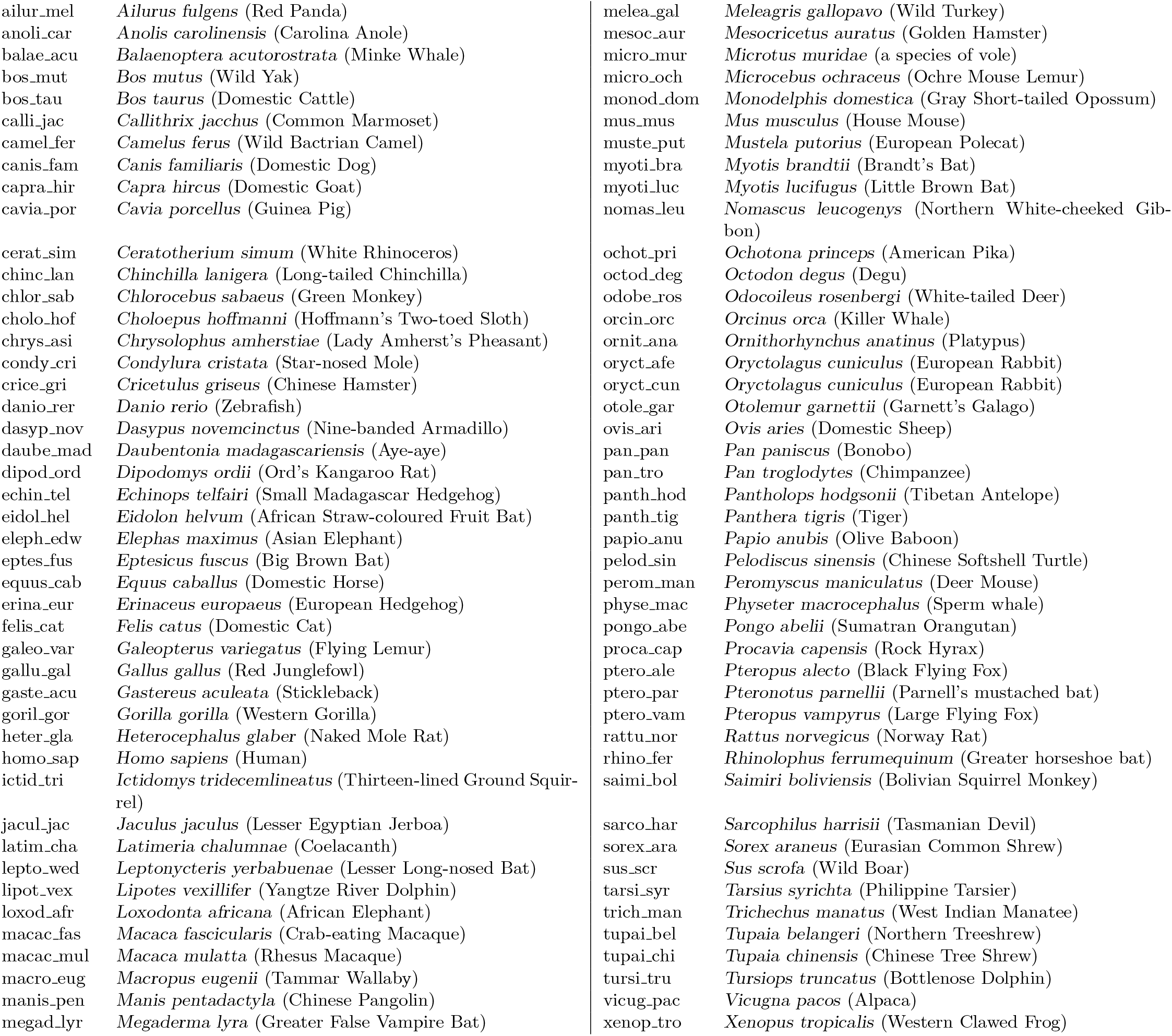
Alphabetical list of Mammal species

**Figure S4.**
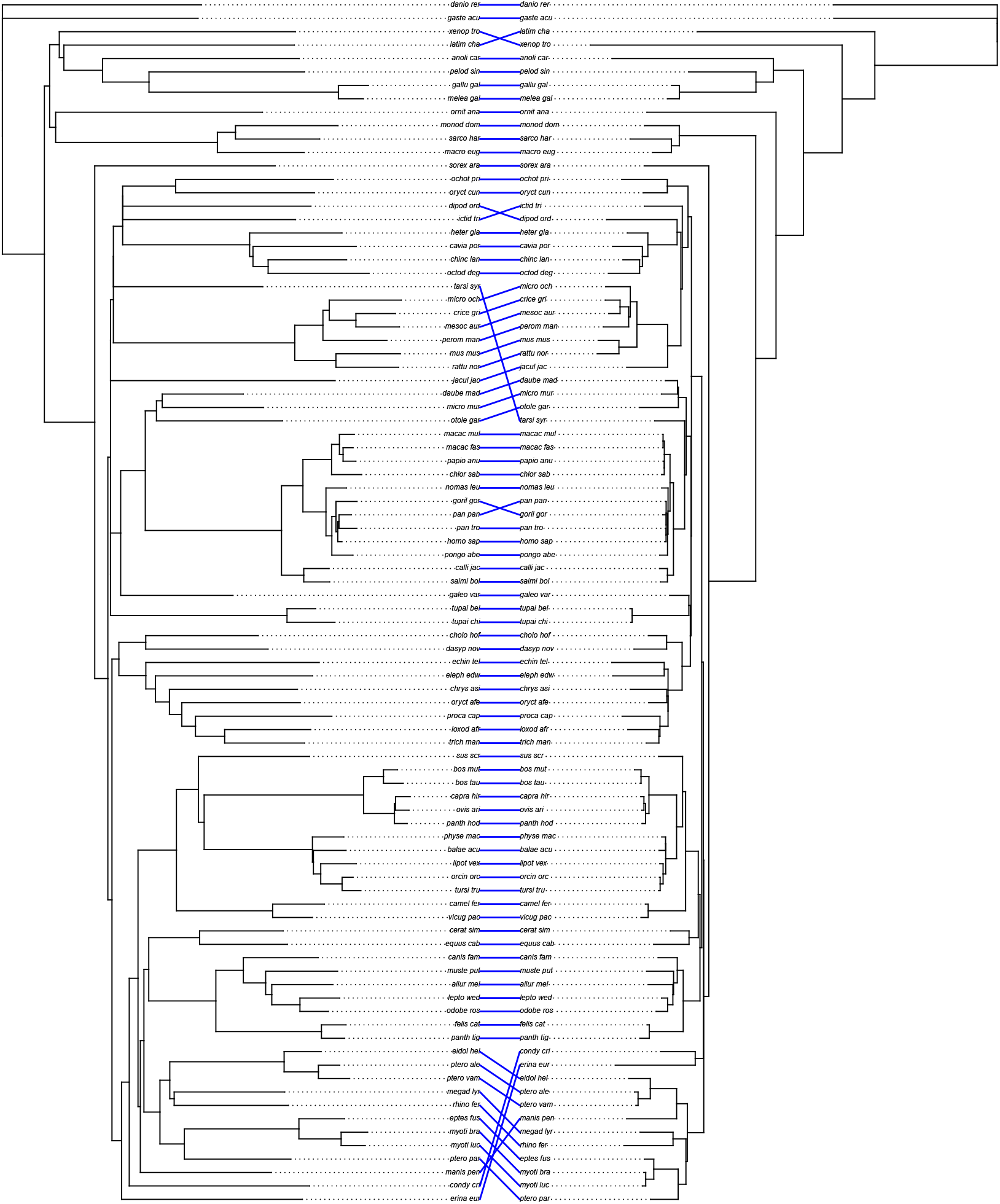
Tanglegram of the mammal dataset comparing the TopicContml tree (left), generated by first removing alignment columns with gaps and then excluding *k*-mers containing ‘N’, to the maximum likelihood tree from Liu et al. (2017) (right). The alphabetical list of the species names in the tree is in Supplement Table S1.

**Figure S5.**
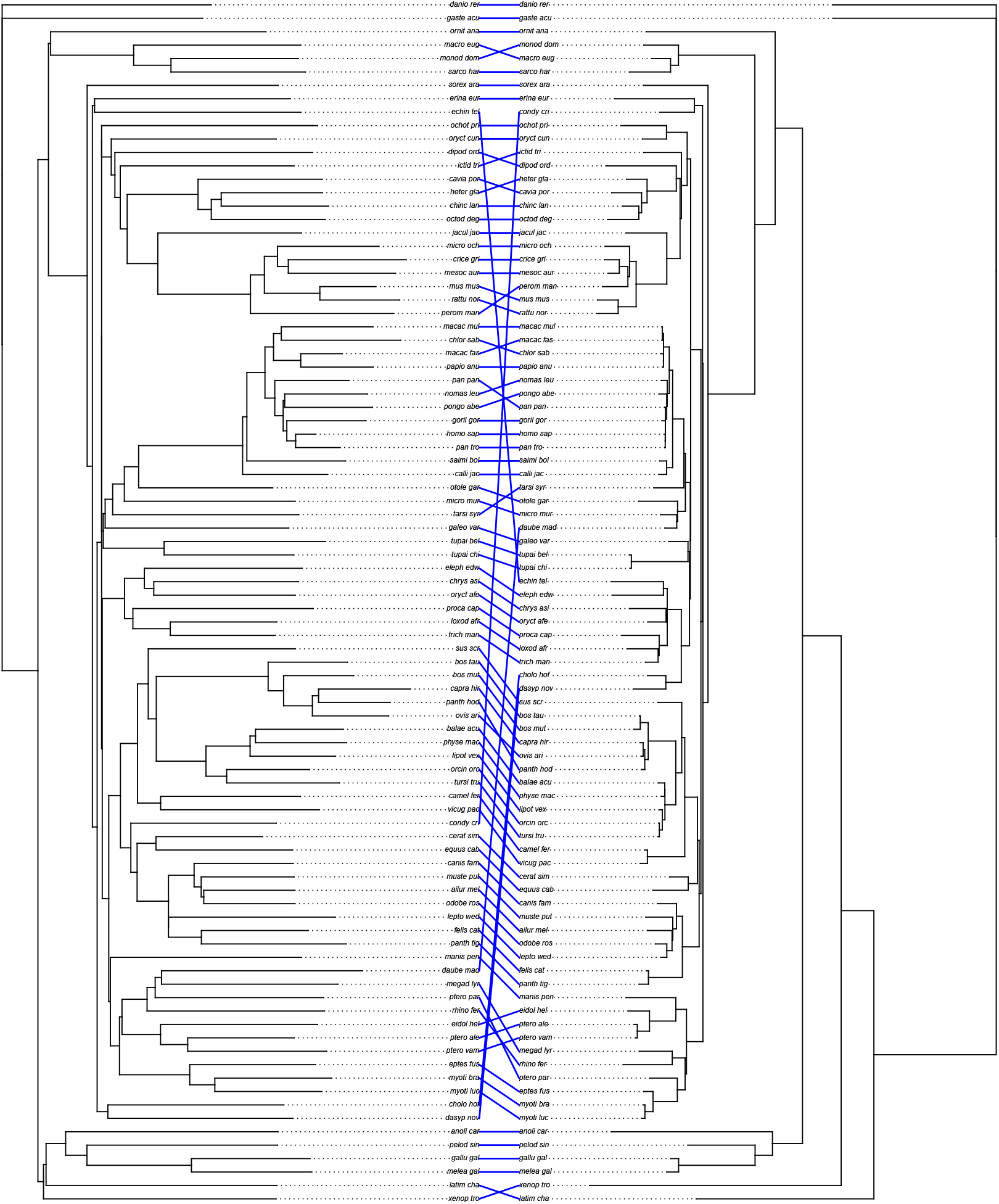
Tanglegram of the mammal dataset comparing the TopicContml tree (left), constructed by removing all gaps from each sequence before excluding *k*-mers containing ‘N’, to the maximum likelihood tree from Liu et al. (2017) (right). The alphabetical list of the species names in the tree is in Supplement Table S1.

**Figure S6.**
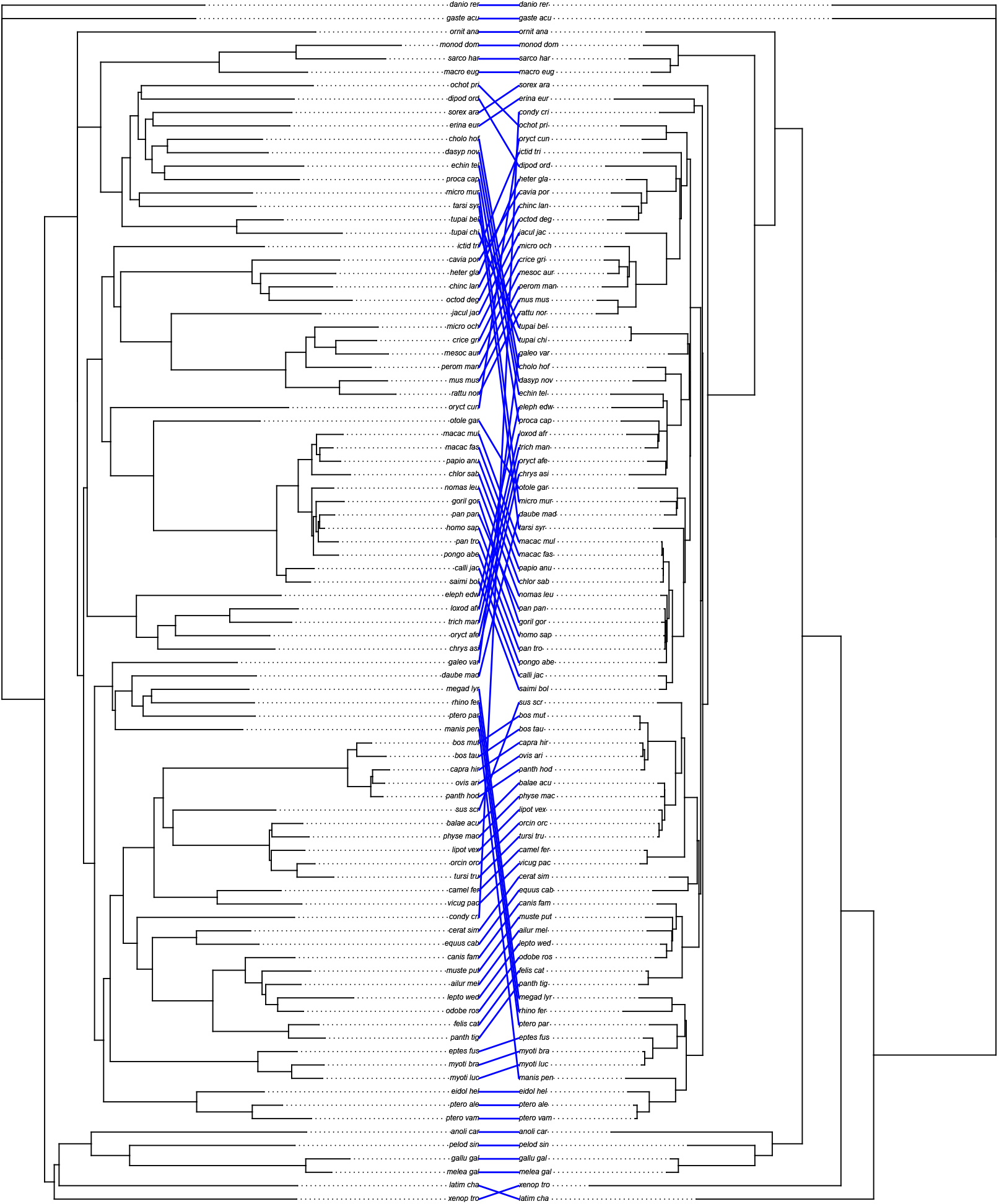
Tanglegram of the mammal dataset comparing the TopicContml tree (left), constructed using aligned sequences with ‘N’ characters retained, to the maximum likelihood tree from Liu et al. (2017) (right). The alphabetical list of the species names in the tree is in Supplement Table S1.

**Figure S7.**
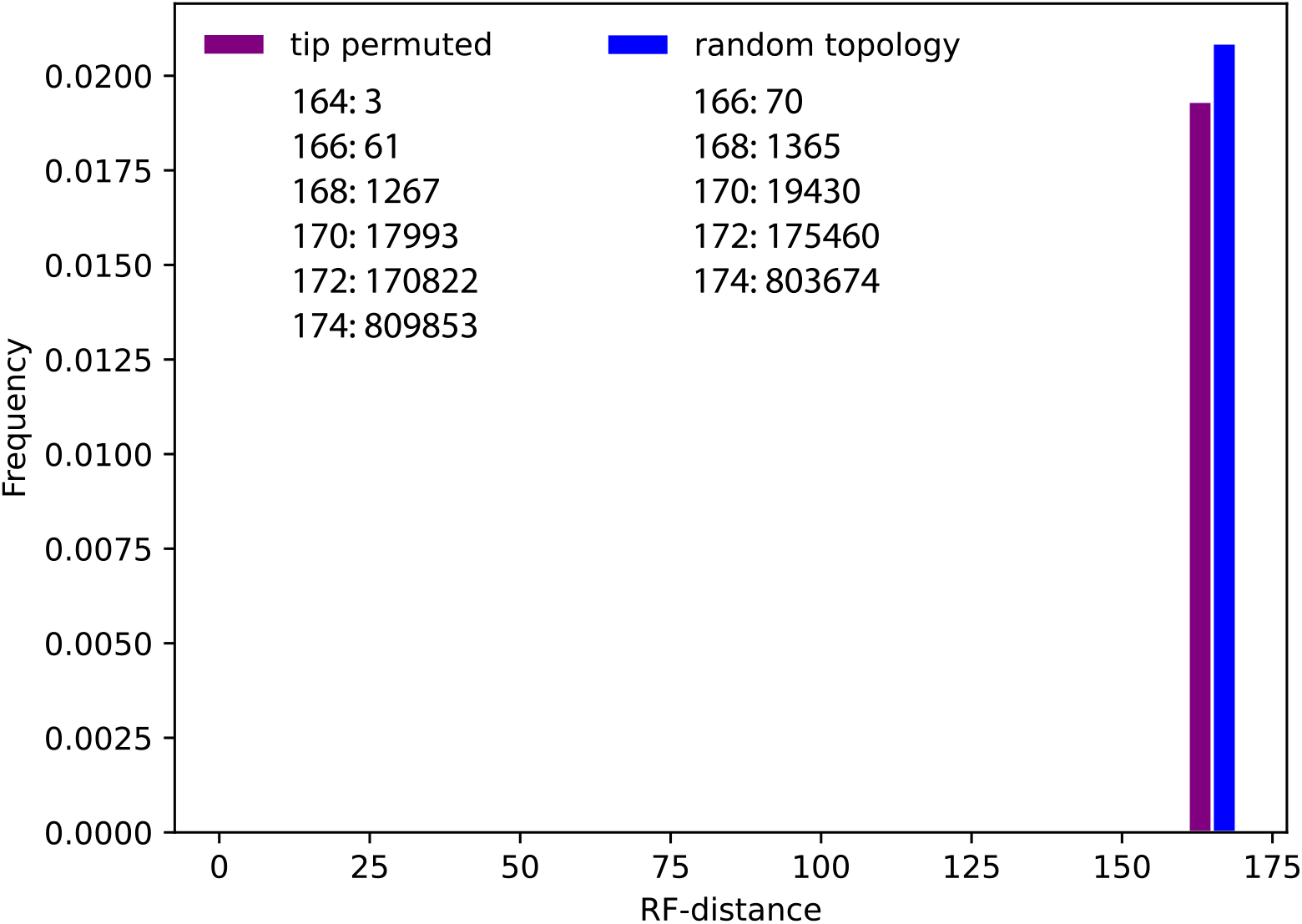
Robinson-Foulds (RF) distance histogram of the maximum likelihood tree of 90 species ‘mammal’ tree (See text) and 1,000,000 random trees with the same number of tips (random topology) and 1,000,000 trees with the true topology but with randomly permuted tip labels. The maximum likelihood tree is at 0. The inserted tables (organized as RF-steps:occurrence) give the exact distributions for each histogram. Our best TopicContml tree was 60 RF-steps from the reference tree.

### PacBio dataset

#### Processing of PacBio Reads

Raw PacBio HiFi reads were downloaded in fastq.gz format from NCBI using the SRA Toolkit (https://hpc.nih.gov/apps/sratoolkit.html#doc) and from the European Nucleotide Archive for the Common Eider. To ensure uniformity of data type and quality, only PacBio HiFi reads were used. Seqkit (Shen et al. 2016) was used to obtain summary statistics for each fastq.gz file, including minimum, maximum and average read lengths. We used seqtk (https://github.com/lh3/seqtk) to subsample 100,000 or 200,000 random reads per species.

**Table S2.**
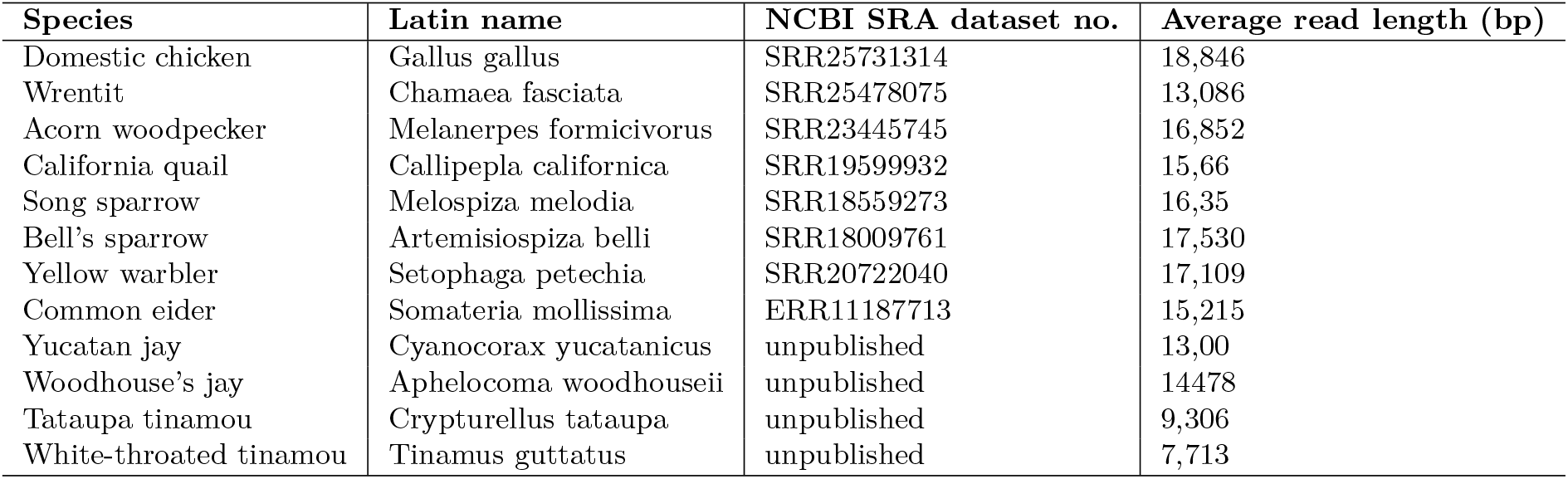
Sources of PacBio long-read sequences from birds for TopicContml analyses.

**Figure S8.**
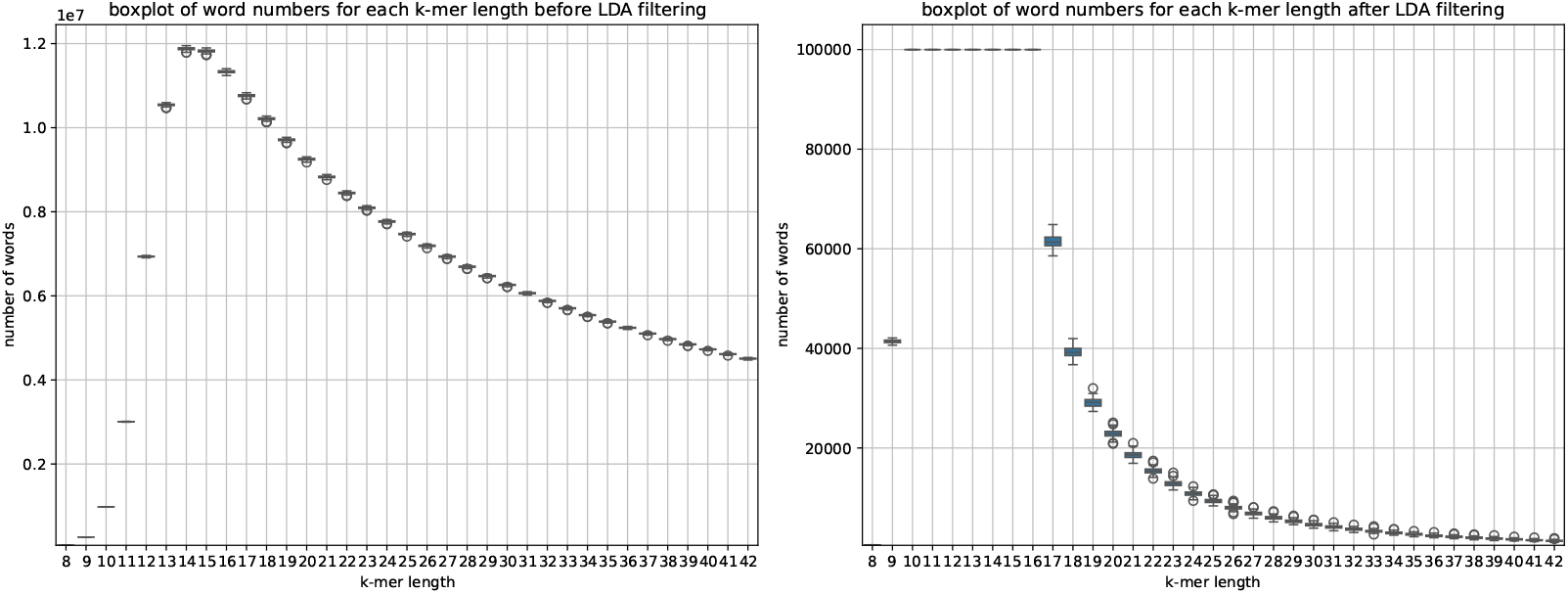
Comparison of word count distributions across loci and *k*-mer lengths ranging from 8 to 42, before and after LDA filtering for the “Unassembled Bird” dataset. The left panel depicts the variation in word numbers prior to filtering, while the right panel demonstrates the impact of LDA in refining and standardizing these distributions. Words (*k*-mers) appearing in fewer than 2 documents or more than 50% of documents are filtered out, retaining only the 100,000 most frequent tokens to optimize memory usage.

**Figure S9.**
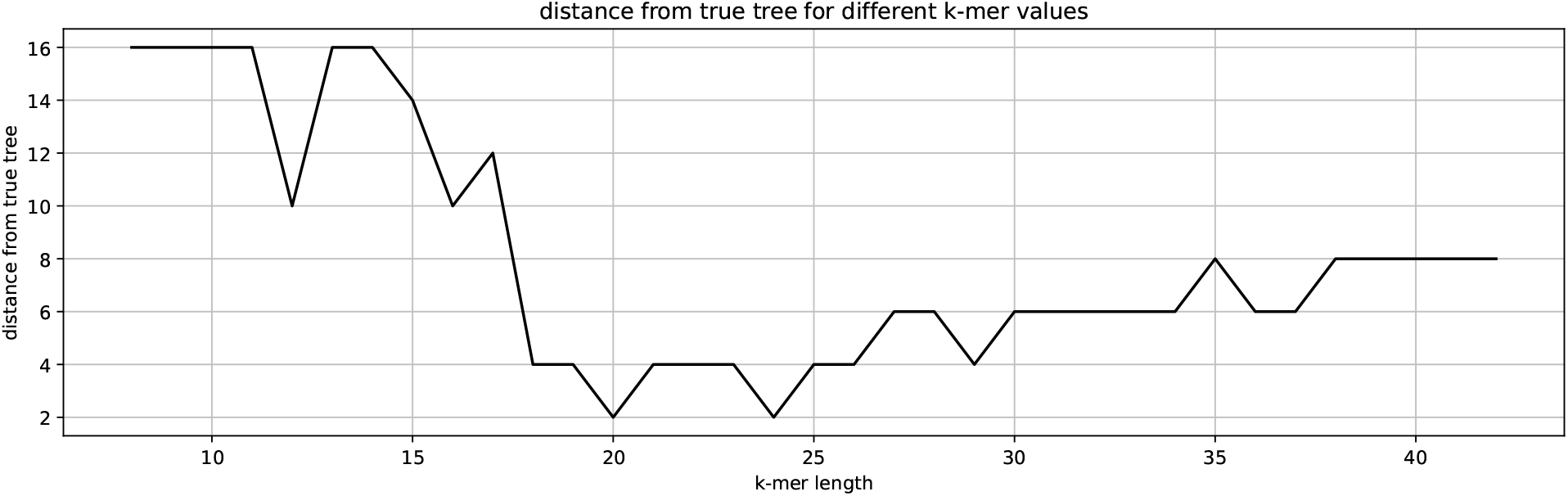
Distance between the true tree and TopicContml -inferred phylogenies across varying *k*-mer lengths for the “Unassembled Bird” dataset. The plot shows the distances of trees generated by TopicContml from the true tree, with the closest agreement observed in the 20-30 *k*-mer length range.

### Runtime for different datasets

**Table S3.**
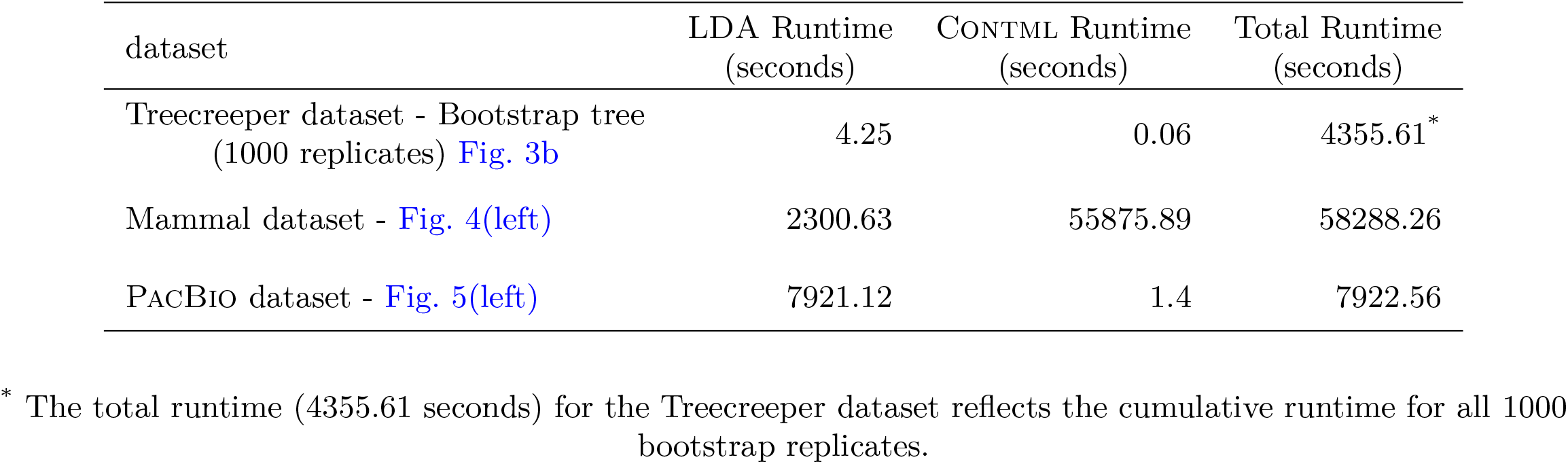
TopicContml runtime for different datasets. The first column lists the dataset names and the corresponding trees generated. The second column shows the runtime for LDA mode (if bootstrapping is applied, it shows approximate LDA time per replicate), part of phase 1. The third column displays the runtime for Contml (if bootstrapping is applied, it shows approximate Contml time per replicate), and the fourth column provides the total elapsed runtime of TopicContml (if bootstrapping is applied, it shows the total elapsed time for all replicates).

